# Efficient multiplex genome editing using CRISPR-Mb3Cas12a in mice

**DOI:** 10.1101/731646

**Authors:** Zhuqing Wang, Yue Wang, Shawn Wang, Andrew J Gorzalski, Hayden McSwiggin, Tian Yu, Kimberly Castaneda-Garcia, Huili Zheng, Wei Yan

## Abstract

Despite many advantages over Cas9, Cas12a has not been widely used in genome editing in mammalian cells largely due to its strict requirement of the TTTV protospacer adjacent motif (PAM) sequence. Here, we report that Mb3Cas12a (*Moraxella bovoculi AAX11_00205*) could edit the genome in murine zygotes independent of TTTV PAM sequences and with minimal on-target mutations and close to 100% editing efficiency when crRNAs of 23nt spacers were used.

**Summary statement:** CRISPR-Mb3Cas12a can target a broader range of sequences in murine zygotes compared to AsCas12a and LbCas12a, and has lower on-target effects than Cas9 and high overall knock-in efficiency.

## Introduction

The rapid advancement of CRISPR-Cas-based genome editing technologies has made gene therapy increasingly promising. However, several obstacles remain, including safety concerns due to both off-target and on-target mutations (Adikusuma et al., 2018; Fu et al., 2013; Hsu et al., 2013; Kosicki et al., 2018; Lee and Kim, 2018) and the requirement of proper PAM sequences for efficient and precise cleavage by the commonly used endonucleases (Komor et al., 2017), e.g., Cas9 and Cas12a/Cpf1. The Cas12a endonuclease has several advantages over Cas9. First, the most commonly used SpCas9 requires NGG PAM sequence, whereas the other widely used AsCas12a and LbCas12a utilize TTTV PAM sequences for efficient genome editing (Cong et al., 2013; Jinek et al., 2012; Wang et al., 2013; Zetsche et al., 2015). Second, Cas12a is guided by a single short CRISPR RNA (crRNA) and can efficiently process its own crRNAs, while Cas9 is directed by dual RNAs consisting of a crRNA and a tracrRNA, and rarely processes its own crRNAs (Cong et al., 2013; Fonfara et al., 2016; Jinek et al., 2012; Zetsche et al., 2017a). Third, Cas12a has much reduced off-target effects compared to SpCas9 due to its irreversible binding to the target region and strong discrimination against the off-target sequences (Kim et al., 2017; Kleinstiver et al., 2016; Strohkendl et al., 2018). Finally, Cas9 has been shown to cause on-target mutations including large deletions and insertions (Adikusuma et al., 2018; Kosicki et al., 2018; Lee and Kim, 2018), whereas Cas12a only generates staggered DNA overhangs, which may lead to much lower rate of on-target mutations due to the so-called preferred microhomology-mediated end joining (MMEJ) repair mechanism (Zetsche et al., 2015). However, practical applications of Cas12a have been severely hindered due, at least in part, to its strict requirement for the TTTV PAM sequence. Although the PAM sequence for FnCas12a has been shown to be YTV (Y stands for C/T and V for A/C/G), the actual editing efficiency of YTV PAM sequences in mammalian cells remains rather low (Tu et al., 2017; Zetsche et al., 2015). Inspired by a recent report suggesting that Mb3Cas12a edits HEK293 cells at a much higher efficiency through TTV PAM sequences compared to AsCas12a and LbCas12a (Zetsche et al., 2017b preprint), we explored whether Mb3Cas12a could be utilized for efficient genome editing and production of knockout/knock-in mice lines.

## Results and Discussion

To determine whether Mb3Cas12a is active in mouse zygotes, we first used one crRNA harboring 20nt direct repeats with 20nt spacer recognizing the TTTV PAM sequence of *Prps1l1* (Table 1, Supplementary Fig. 1A, D), a testis-specific gene dispensable for spermatogenesis (Wang et al., 2018). One out of six founders obtained has an indel (insertion or deletion) (16.7%) (Table 1, Supplementary Fig. 1A, D), suggesting that Mb3Cas12a indeed works in mouse zygotes. Since AsCas12a and LbCas12a have the ability to process their own crRNAs (Fonfara et al., 2016; Zetsche et al., 2017a), we next tested whether Mb3Cas12a could do the same. We designed one crRNA harboring two 20nt spacers recognizing two TTTV PAM sequences at *Saraf* locus (Table 1, Supplementary Fig. 1B, E). The 20nt spacers were separated by 20nt direct repeats. One out of three founders was edited by spacer 1 (33.3%) (Table 1, Supplementary Fig. 1B, E), suggesting that Mb3Cas12a can indeed process its own crRNAs to edit a specific locus in mouse zygotes. We then determined whether Mb3Cas12a could utilize the TTV PAM sequence in mouse zygotes. We designed one crRNA harboring two 20nt spacers recognizing two TTV PAM sequences of the same *Saraf* locus (Table 1, Supplementary Fig. 1B, E). One out of four founders was edited with the spacer 2 (25%) (Table 1, Supplementary Fig. 1B, E), indicating that Mb3Cas12a can indeed target genomic DNA with TTV PAM sequences. Since the orientation of crRNAs has no significant effect on genome editing efficiency (Zetsche et al., 2017a), we compared the efficiency of Mb3Cas12a in editing DNA harboring TTTV and TTV PAM sequences by utilizing one single crRNA containing two spacers recognizing TTTV and TTV PAM sequences, respectively, in the *Mrvi1* locus (Table 1, Supplementary Fig. 1C, F). Out of 5 founders, one was edited with the spacer recognizing the TTTV PAM sequence (20%), but none from the TTV PAM sequence (Table 1, Supplementary Fig. 1C, F), suggesting that Mb3Cas12a prefers the spacer targeting the TTTV PAM sequence.

**Table 1.**
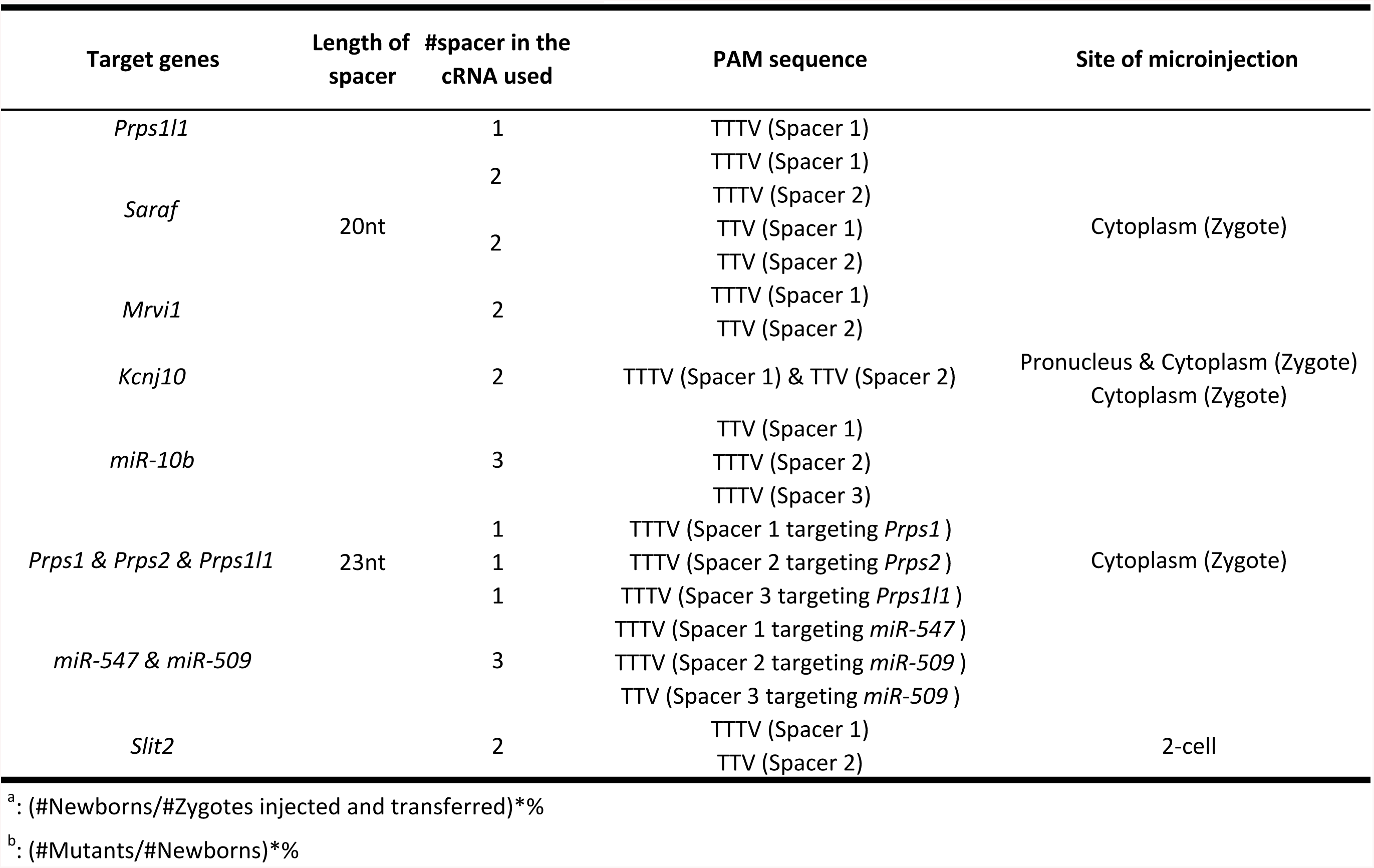
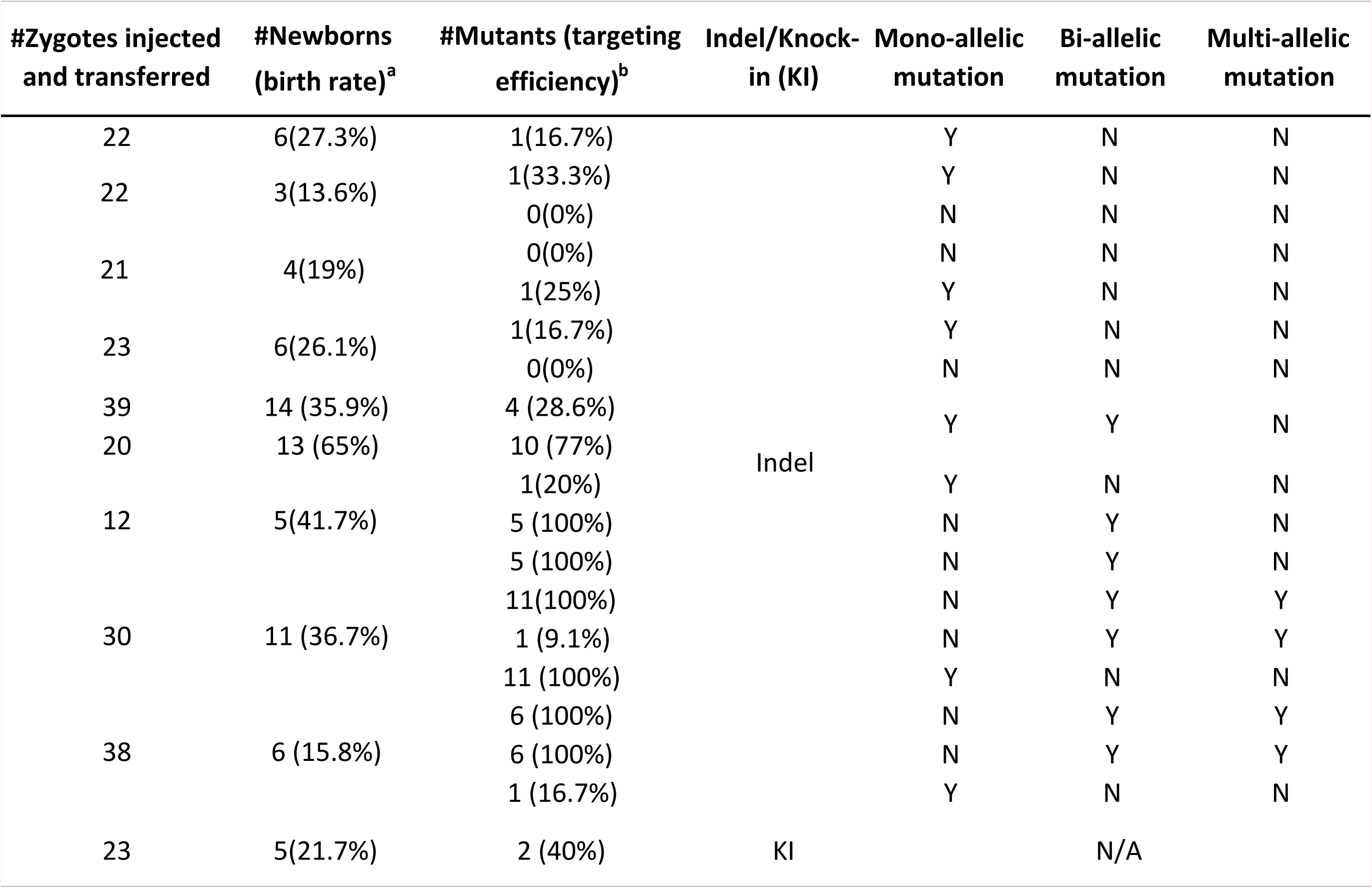
Editing efficiency of Mb3Cpf1 in murine zygotes.

As the length of crRNAs has been shown to affect the efficiency of Cas12a-mediated genome editing (Tu et al., 2017; Zetsche et al., 2015), we further tested whether the editing efficiency can be improved by optimizing the crRNA structure. We first tested effects of the length of crRNAs on genome editing in *Dnmt1* locus in HEK293 cells (Fig. 1A, B). No Mb3Cas12a activity was detected when a 17nt spacer was used, whereas the highest Mb3Cas12a activities comparable to AsCas12a and LbCas12a were observed in the crRNAs with 23nt spacer (Fig. 1B). Similar results have been reported for AsCas12a and LbCas12a, but not for FnCas12a, which appears to use 21nt crRNAs more efficiently (Tu et al., 2017). To determine the potential effects of microinjection (cytoplasmic vs. pronuclear) methods on targeting efficiency, we injected Mb3Cas12a mRNA and one crRNA, harboring two spacers (one recognizing TTTV PAM sequence, and one TTV) targeting *Kcnj10* locus, into either cytoplasm only or both pronucleus and cytoplasm. Interestingly, the cytoplasmic injection appeared to have a higher targeting efficiency (77%, n=20) than the pronuclear and cytoplasmic injection (28.6%, n=39) (Table 1, Fig. 1C, D). Therefore, we used 23nt spacer and cytoplasmic microinjection to generate the following indels in mice (Fig. 2A).

**Figure 1.**
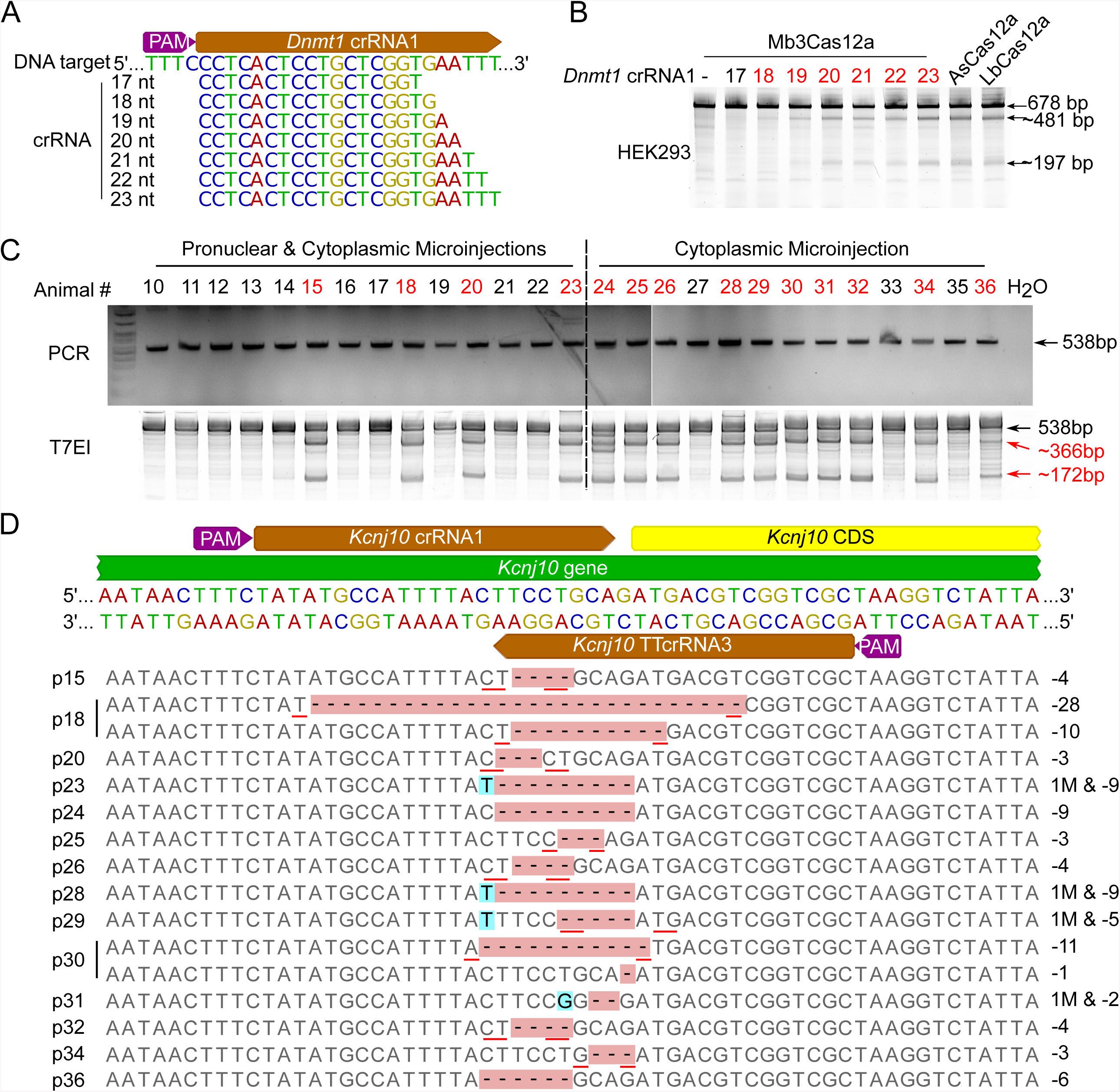
Optimization of Mb3Cas12a-based genome editing efficiency by adjusting the crRNA length and microinjection sites. A. Sequences of crRNAs targeting *Dnmt1* locus. crRNAs with one 20nt direct repeat and spacers of various lengths recognizing TTTV PAM sequences were used for targeting *Dnmt1* in HEK293 cells. B. T7EI assay results of Mb3Cas12a-edited *Dnmt1* locus in HEK293 cells. C. PCR and PCR-T7EI to identify the efficiency of Mb3Cas12a-based genome editing in *Kcnj10* locus by either pronuclear and cytoplasmic microinjection or cytoplasmic microinjection only in murine zygotes. D. crRNAs used in targeting *Kcnj10* locus (upper panel) and Sanger sequencing results of Mb3Cas12a-based genome editing in *Kcnj10* locus in murine zygotes (lower panel). Red underlines represent microhomology (MH) sequences.

**Figure 2.**
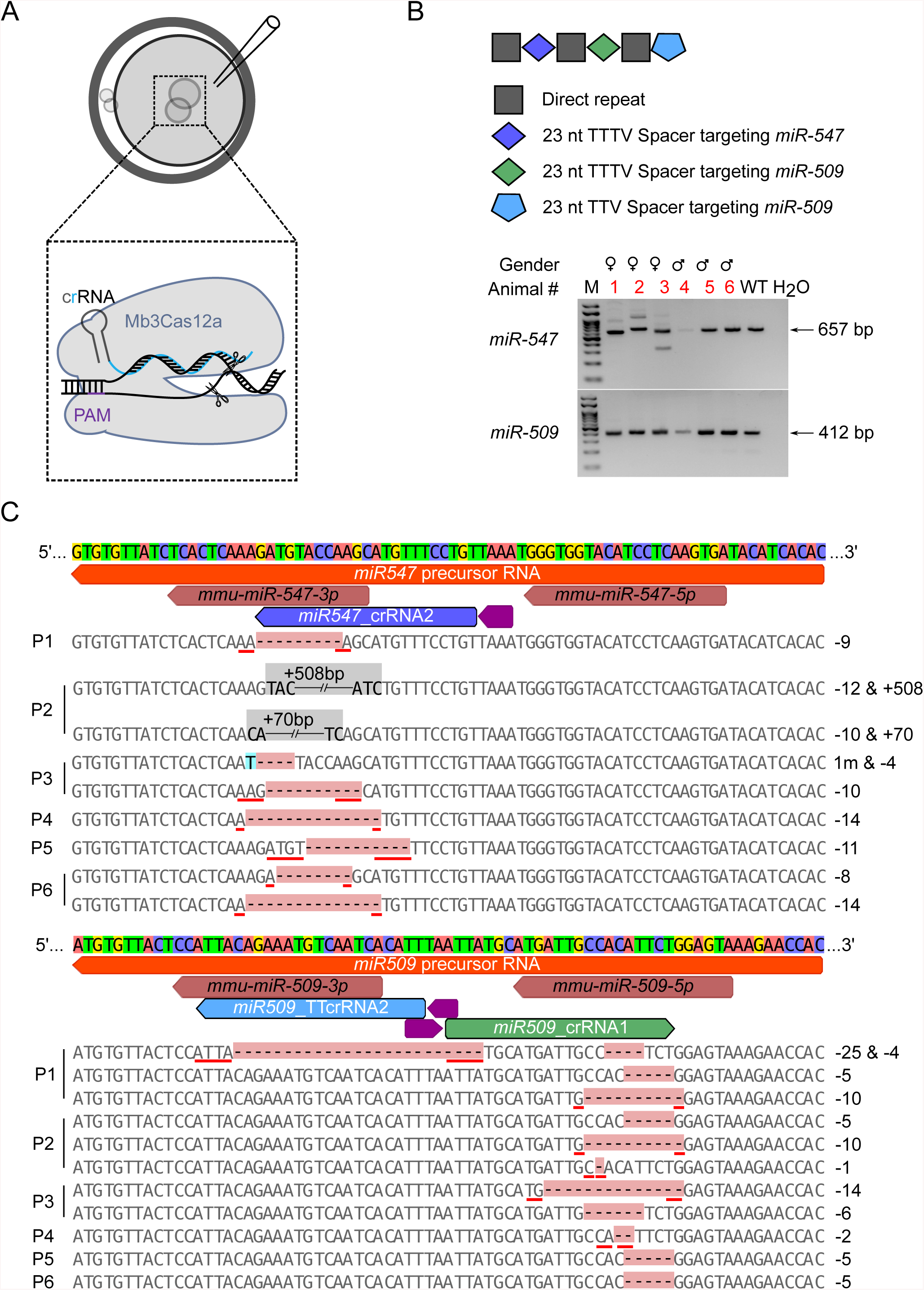
Multiplex targeting efficiency of Mb3Cas12a in *miR-547* & *miR-509* loci with TTV PAM sequence in mice. A. Schematics showing the strategy used for generating indels in mouse zygotes. Mb3Cas12a mRNA and 23nt crRNA are microinjected into the cytoplasm of mouse zygotes. B. PCR genotyping results of Mb3Cas12a-edited *miR-547 & miR-509* founders. One crRNA with one 23nt spacer targeting *miR-547* with a TTTV PAM sequence and two 23nt spacers targeting *miR-509* with one TTTV PAM and one TTV PAM sequences was used to target *miR-547 & miR-509* loci. C. Sanger sequencing results of Mb3Cas12a-edited pups #1, #2, #3, #4, #5 and #6 in *miR-547 & miR-509* loci. Red underlines represent microhomology (MH) sequences, characters in grey background indicate large insertions.

To explore whether Mb3Cas12a could target multiple loci simultaneously, we used one crRNA harboring three 23nt spacers recognizing TTTV PAM sequencing targeting the *Prps* family, i.e. *Prps1, Prps2* and *Prps1l1*. Eleven founders obtained (100%) were edited by both the *Prps1* and *Prps1l1* crRNAs, whereas one (9.1%) was edited only by the *Prps2* crRNA (Table 1, Supplementary Fig. 2A), which may reflect a lower targeting efficiency of this particular crRNA. Next, we used a crRNA containing three 23nt spacers separated by 20nt direct repeats to target *miR-10b* (Table 1, Supplementary Fig. 2B, C). Two of the spacers (spacers 2 and 3) were designed to recognize a TTTV PAM sequence, whereas the other one targets a TTV PAM sequence (spacer 1) at *miR-10b* locus. All the five founders were edited by the two spacers targeting the TTTV PAM sequence (100%), and one of them was edited by the one recognizing the TTV PAM sequence (20%) (Table 1, Supplementary Fig. 2B, C). Similar results were also obtained when we used one crRNA containing one 23nt spacer targeting *miR-547* with a TTTV PAM sequence and two 23nt spacers targeting *miR-509* with one TTTV PAM and one TTV PAM. All six founders were edited by the two spacers targeting the TTTV PAM sequence (100%), and one of them was edited by the one recognizing the TTV PAM sequence (16.7%) (Table 1, Fig. 2B, C). Moreover, although the 23nt spacers recognizing TTTV PAM sequence often led to bi- or multi-allelic targeting and TTV PAM sequence tended to yield mono-allelic targeting, the 20nt spacers appeared to cause mostly mono-allelic mutations (Table 1, Fig. 1D, 2C and Supplementary Fig. 1, 2). Recent reports have shown that Cas9 with one single gRNA tends to induce large indels in genomic DNA in mouse embryonic stem (ES) cells, progenitor cells and zygotes (Adikusuma et al., 2018; Kosicki et al., 2018), whereas Cas9 with two gRNAs causes large deletions within the two flanking gRNA-targeting sites (Wang et al., 2018). The incidences of large deletions induced by Cas9 with one single gRNA were 35.7%, 36.5%, and 45% in mouse ES cells, progenitor cells, and zygotes, respectively, whereas the incidence of large insertions was 26.3% in mouse ES cells (Adikusuma et al., 2018; Kosicki et al., 2018). Interestingly, unlike Cas9, Mb3Cpf-based genome editing predominantly generates indels, containing microhomology (MH) sequences flanking the cleavage sites with one or more spacers within a single crRNA. Among all 42 pups derived from Mb3Cas12a-based editing, only 3 contained large insertions (7.1%) induced by 3 spacers in the crRNA, whereas the rest displayed different alleles with either two or more small mutations or one mutant plus one wild-type alleles (Table 1, Fig. 1D, 2C and Supplementary Fig. 1, 2). Based on RepeatMasker, two large insertions correspond to ERVL (endogenous retroviruses type-L) and ERVL-MaLR (mammalian apparent LTR retrotransposon), respectively, similar phenomenon has been observed in SpCas9 induced double strand breaks (DSBs) (Ono et al., 2015), suggesting these retrotransposons may hijack the DSBs induced by SpCas9 or Mb3Cas12a. None of the Mb3Cas12a-edited alleles contained large deletions, whereas they are commonly seen in Cas9-editted genes. These results suggest that MMEJ repair mechanism is preferentially adopted in fixing the staggered DNA ends, which may account for the minimal on-target effects in Mb3Cas12a-based genome editing.

Given that two-cell homologous recombination (2C-HR)-CRISPR, in which Cas9 was tethered with monomeric streptavidin (mSA) that could bind to biotinylated DNA donor template, showed a higher knock-in (KI) efficiency (Gu et al., 2018), we tested whether Mb3Cas12a-mSA could do the same in generating KI mice (Fig. 3A, B). We microinjected 2-cell embryos with Mb3Cas12a-mSA mRNA, crRNA targeting *Slit2* locus, and biotinylated DNA donor template containing a BamHI restriction enzyme cutting site (Fig. 3A). Among 5 founders, 2 (40%) have the knock-in alleles, indicating the Mb3Cas12a-mSA indeed can generate KI mice efficiently (Fig. 3B, 3C). During preparation of our manuscript, one study reporting that HkCas12a can target YTV and TYYN PAM sequences in human cell lines was published (Teng et al., 2019). However, it remains unknown whether HkCas12a works in murine zygotes and what its efficiency is. In summary, our data demonstrate that Mb3Cas12a can edit the murine genome independent of TTTV PAM sequence and with minimal on-target mutations and very high targeting efficiency. Mb3Cas12a-mediated genome editing expands the toolkit for efficient production of mutant mouse lines.

**Figure. 3.**
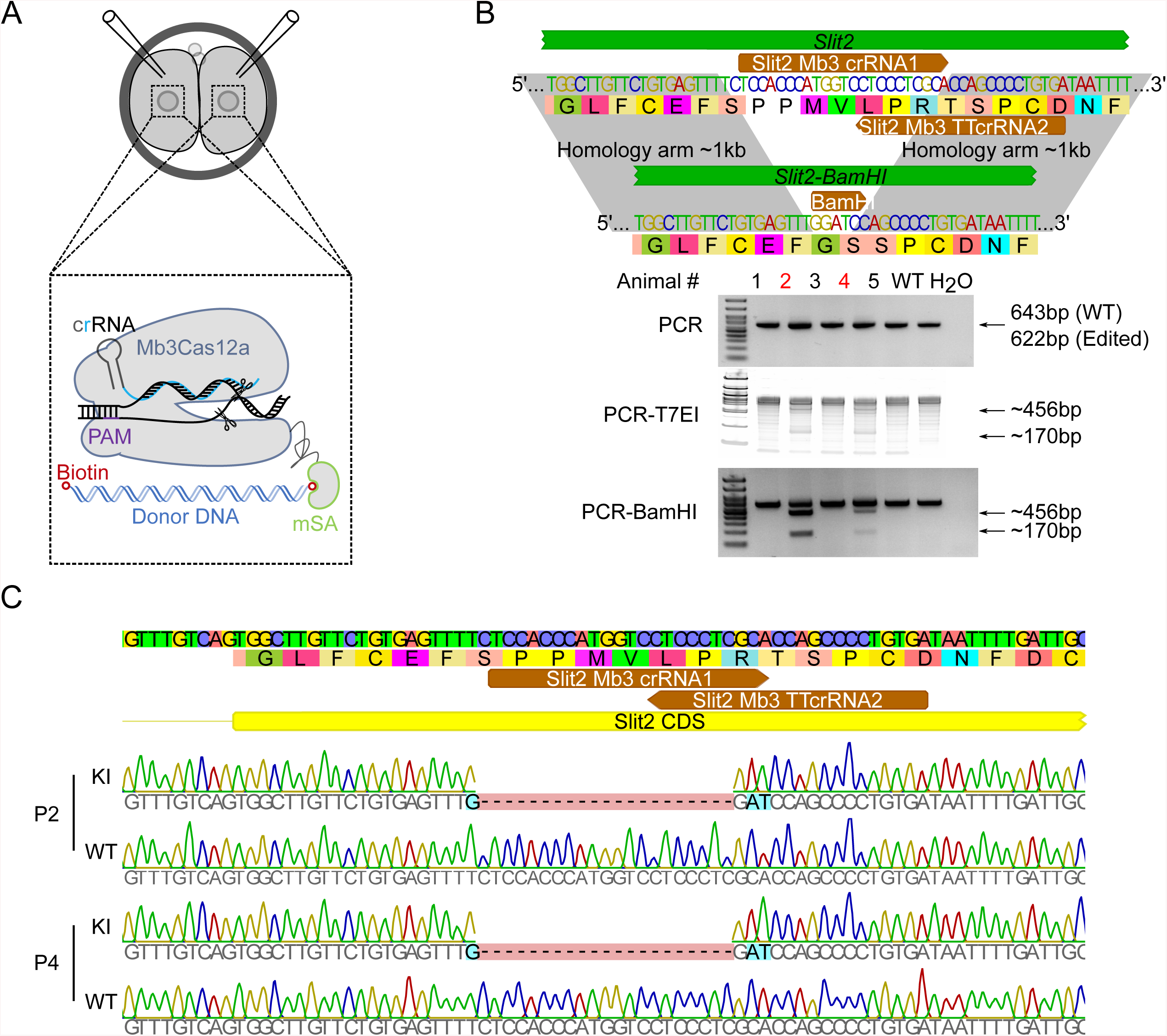
Generation of knock-in (KI) in *Slit2* locus using Mb3Cas12a-mSA in mouse 2-cell embryos. A. Schematics showing the strategy used for generating KI in 2-cell mouse embryos. Mb3Cas12a mRNA, crRNA and biotinylated donor DNA template are microinjected into 2-cell mouse embryos. B. Efficiency of Mb3Cas12a-mSA-mediated KI in *Slit2* locus in mice. One crRNA harboring two 23nt spacers recognizing one TTTV PAM and one TTV PAM sequence was used to target *Slit2* locus. Upper panel, strategy used for generating *Slit2-BamHI* KI. Colored characters represent DNA sequence, black characters in colored background indicate corresponding amino acids. Lower panel, PCR, PCR-T7EI (T7 endonuclease I assay) and PCR-BamHI digestion show the KI efficiency. C. Sanger sequencing results of Mb3Cas12a-mSA-mediated KI pups #2, and #4 in *Slit2* locus.

## Materials and Methods

### Plasmids construction

To prepare pcDNA3.1-Mb3Cas12a-mSA plasmid, monomeric streptavidin (mSA) DNA fragments amplified from PCS2+Cas9-mSA plasmid (Cat. 103882, Addgene, Watertown, MA) were inserted into the pY117 plasmid (pcDNA3.1-huMb3Cpf1) (Cat. 92293, Addgene) *via* BamHI (Cat. R0136S, NEB, Ipswich, MA) and EcoRI (Cat. R3101S, NEB) restriction sites.

For the pUC-Slit2-BamHI plasmid, two homology arms (∼1kb) flanking the crRNAs cutting sites of *Slit2* locus and pUC empty vector were amplified by Q5® Hot Start High-Fidelity 2X Master Mix (Cat. M0494S, NEB) from mouse tail genomic DNA and pX330 plasmid (Cat. 42230, Addgene), respectively. After purification with Ampure beads, these three DNA fragments were assembled with NEBuilder® HiFi DNA Assembly Master Mix (Cat. E2621L, NEB). BamHI restriction site was introduced between the two homology arms during the PCR amplification. The primers used for plasmids construction are listed in **Supplemental Table S1**.

### Generation of Mb3Cas12a and Mb3Cas12a-mSA mRNA, crRNAs, and donor DNA template

To synthesize Mb3Cas12a and Mb3Cas12a-mSA mRNAs, the pY117 plasmid (pcDNA3.1-huMb3Cpf1) (Cat. 92293, Addgene) and pcDNA3.1-Mb3Cas12a-mSA were digested with EcoR I (Cat. R3101S, NEB) overnight at 37°C, followed by purification with Ampure beads and mRNA synthesis with the HiScribe™ T7 ARCA mRNA Kit (Cat. E2065S, NEB). Then the *in vitro* transcribed mRNAs were treated with DNase I (NEB, Cat. M0303S) to remove the plasmid DNA template, followed by poly(A) tailing using E. coli poly(A) polymerase (Cat. M0276S, NEB). The poly(A)-tailed Mb3Cas12a mRNAs were purified using the RNA Clean & Concentrator™-5 (Cat. R1016, Zymo Research, Irvine, CA) and eluted in a Tris-EDTA solution (Cat.11-01-02-02, IDT, Coralville, IA).

crRNAs were designed using Benchling (https://benchling.com/). DNA oligos for making each crRNA were synthesized in the IDT Inc. and listed in **Supplemental Table S1**. To prepare crRNAs for microinjection, the T7 first strand primer and antisense oligos specific for each crRNA were mixed in 1X T4 DNA ligase buffer and heated to 95°C for 5 minutes, and then allowed to cool down to room temperature on the bench. The annealed oligos were used as the templates for *in vitro* transcription (IVT) using the HiScribe™ T7 High Yield RNA Synthesis Kit (Cat. E2040S, NEB). After IVT, crRNAs were purified using the RNA Clean & Concentrator™-5 (Cat. R1016, Zymo Research) and eluted in Tris-EDTA solution (Cat.11-01-02-02, IDT).

To prepare crRNAs for transfection of HEK293 cells, PCR products corresponding to each crRNA were amplified with U6 forward primer and corresponding antisense oligos (as listed in **Supplemental Table S1)** from the pX330 plasmid (Cat. 42230, Addgene). After digestion with DpnI (Cat. R0176S, NEB), the PCR products were purified using Ampure beads.

The biotinylated donor DNA template was amplified from the pUC-Slit2-BamHI plasmid with biotinylated primers (as listed in **Supplemental Table S1)**, followed by DpnI digestion to remove the plasmid and purification with Ampure beads.

### HEK293 cells Transfection

HEK293 cells were co-transfected with 400ng of pY117 (pcDNA3.1-huMb3Cpf1) (Cat. 92293, Addgene) and 100ng of crRNA PCR product using Lipofectamine 2000 (Cat. 11668, Thermo Fisher Scientific, Waltham, MA) in a 24 well cell culture plate (Cat. 3524, Corning, Corning, NY). After 48h, cells were collected for analyses.

### Animal use and generation of knockout (KO) and knock-in (KI) mice

The animal protocol for this study was approved by the Institutional Animal Care and Use Committee (IACUC) of the University of Nevada, Reno (protocol number 00494). All mice were housed and maintained under specific pathogen free conditions with a temperature- and humidity-controlled animal facility in the Department of Lab Animal Medicine, University of Nevada, Reno. Generation of KO and KI mice were performed as previously described with modifications (Gu et al., 2018; Wang et al., 2018; Wang et al., 2019). Briefly, 4-6 weeks of FVB/NJ or C57BL/6J female mice were super-ovulated and mated with C57BL/6J stud males; zygotes and 2-cell stage embryos were collected from the oviducts for KO and KI, respectively. For KO, Mb3Cas12a mRNA (200ng/μl) and crRNA (100 ng/μl) were mixed and injected into the cytoplasm or pronucleus of the zygotes in M2 medium (Cat. MR-051-F, Millipore, Burlington, MA). For KI, Mb3Cas12a-mSA mRNA (75ng/μl), crRNA (50 ng/μl) and biotinylated donor DNA template (20ng/μl) were mixed and injected into the cytoplasm or pronucleus of the 2-cell embryos in M2 medium. After injection, all embryos were cultured for 1h in KSOM+AA medium (Cat. MR-121-D, Millipore) at 37°C under 5% CO_2_ in air before being transferred into 7-10 week-old female CD1 recipients.

### Mouse genotyping, T7EI and Sanger sequencing

Mouse genotyping was performed as previously described (Wang et al., 2018; Wang et al., 2019). Briefly, mouse tail or ear snips were lysed in a lysis buffer (40mM NaOH, 0.2mM EDTA) for 1h at 95°C, followed by neutralization using the same volume of neutralizing buffer (40mM Tris-HCl). PCR reactions were conducted using Platinum™ SuperFi™ Green PCR Master Mix (Cat. 12359010, Thermo Fisher Scientific). T7EI (Cat. M0302L, NEB) assay was followed to detect the mutations. The positive samples were proceeded with A tailing using GoTaq® Green Master Mix (Cat. M7123, Promega, Madison, WI) for 5min at 95 °C, followed by 15min at 72 °C. The A-tailed PCR products were then ligated to pGEM®-T Easy Vector using pGEM®-T Easy Vector Systems (Cat. A1360, Promega). Positive colonies were selected for Sanger sequencing. Data was analyzed using Geneious software. The primers used for genotyping are listed in **Supplemental Table S1**.

### MiSeq library construction and analysis

DNA fragment of *Prps1, Prps2*, and *Prps1l1* were amplified using Platinum™ SuperFi™ Green PCR Master Mix (Cat. 12359010, Thermo Fisher Scientific) from lysis of mouse tail or ear snips. The PCR products were then tagged using Nextera XT DNA Library Preparation Kit (Cat. 15032354, Illumina, San Diego, CA) and indexed using Nextera XT Index Kit (Cat. 15055294, Illumina). DNA library was sequenced using MiSeq Reagent Kit v2 (500-cycles) (Cat. MS-102-2003, Illumina). Data was analyzed using Geneious software. The primers used for *Prps1, Prps2*, and *Prps1l1* are listed in **Supplemental Table S1**.

## Acknowledgements

Not applicable.

## Data availability

The datasets generated and/or analyzed during the current study are available in the Sequence Read Archive (SRA), https://www.ncbi.nlm.nih.gov/sra/PRJNA556550

## Authors’ contributions

Z. W. and W. Y. conceived and designed the research. Z. W., Y. W., S. W., A. J G., H. M., T. Y., K. C-G., and H.Z. performed bench experiments. Z. W. analyzed data. Z. W. and W. Y. wrote the manuscript. All reviewed and agreed with the contents of the manuscript.

---

**Supplementary Figure 1.**
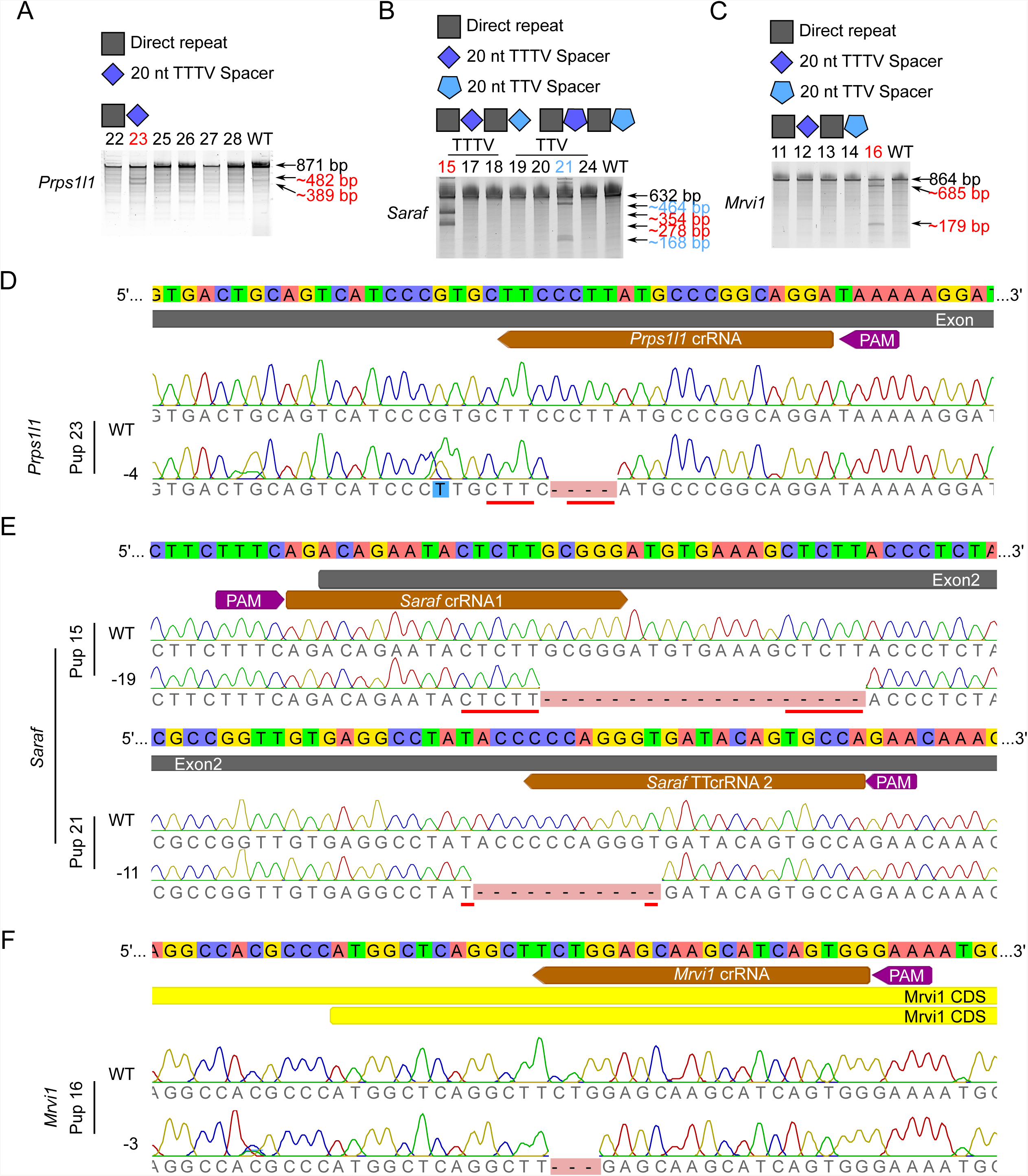
Efficiency of Mb3Cas12a-based genome editing using 20nt spacers targeting *Prps1l1, Saraf* and *Mrvi1* loci in mice. A. T7 endonuclease I (T7EI) assay results of Mb3Cas12a-edited *Prps1l1* locus. One crRNA with one 20nt direct repeat and one 20nt spacer recognizing a TTTV PAM sequence was used for targeting *Prps1l1*. B. T7EI assay results of Mb3Cas12a-edited *Saraf* locus. Two crRNAs harboring two 20nt spacers either recognizing two TTTV or two TTV PAM sequences were used to target *Saraf* locus. The expected bands corresponding to the pups after T7EI assays are indicated with the same color. C. T7EI assay results of Mb3Cas12a-edited *Mrvi1* locus. One single crRNA containing two spacers recognizing both TTTV and TTV PAM sequences was used to target *Mrvi1* locus. D. Sanger sequencing results of Mb3Cas12a-edited pup #23 in *Prps1l1* locus. E. Sanger sequencing results of Mb3Cas12a-edited pups #15 and #21 in *Saraf* locus. F. Sanger sequencing results of Mb3Cas12a-edited pup #16 in *Mrvi1* locus. Red underlines represent microhomology (MH) sequences.

**Supplementary Figure 2.**
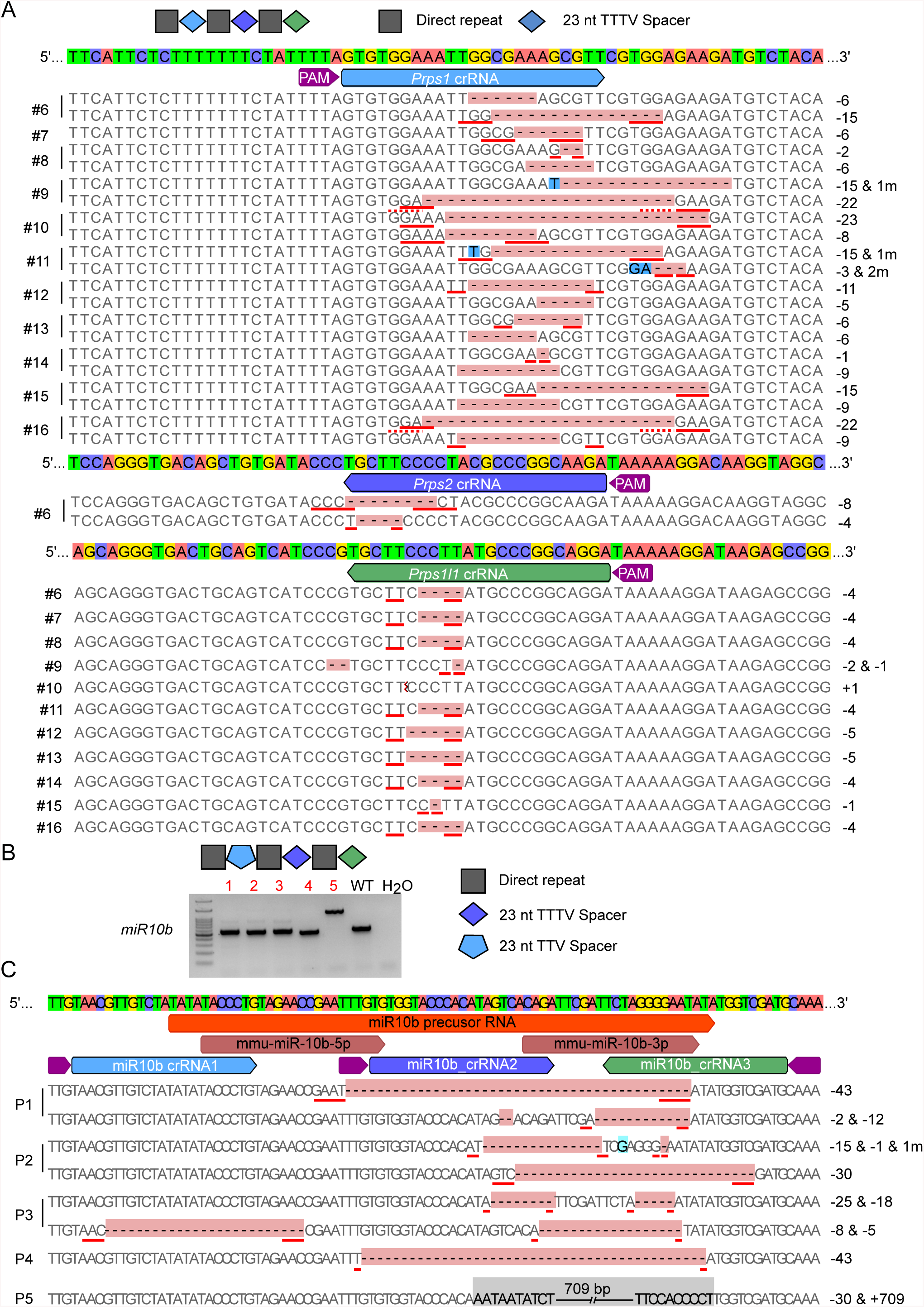
Multiplex targeting efficiency of Mb3Cas12a in *Prps* family and *miR-10b* loci in mice. A. MiSeq results of Mb3Cas12a targeting efficiency in *Prps1, Prps2*, and *Prps1l1* with one crRNA, which contains three 20nt direct repeats and three 23nt spacers targeting *Prps1, Prps2* and *Prps1l1* with TTTV PAM sequences. Red underlines represent microhomology (MH) sequences. B. PCR genotyping results of Mb3Cas12a-edited *miR-10b* founders. One crRNA with three 20nt direct repeats and three 23nt spacers recognizing two TTTV PAM and one TTV PAM sequences was used to target *miR-10b* locus. C. Sanger sequencing results of Mb3Cas12a-edited pups #1, #2, #3, #4 and #5 in *miR-10b* locus.

**Supplemental Table S1.**
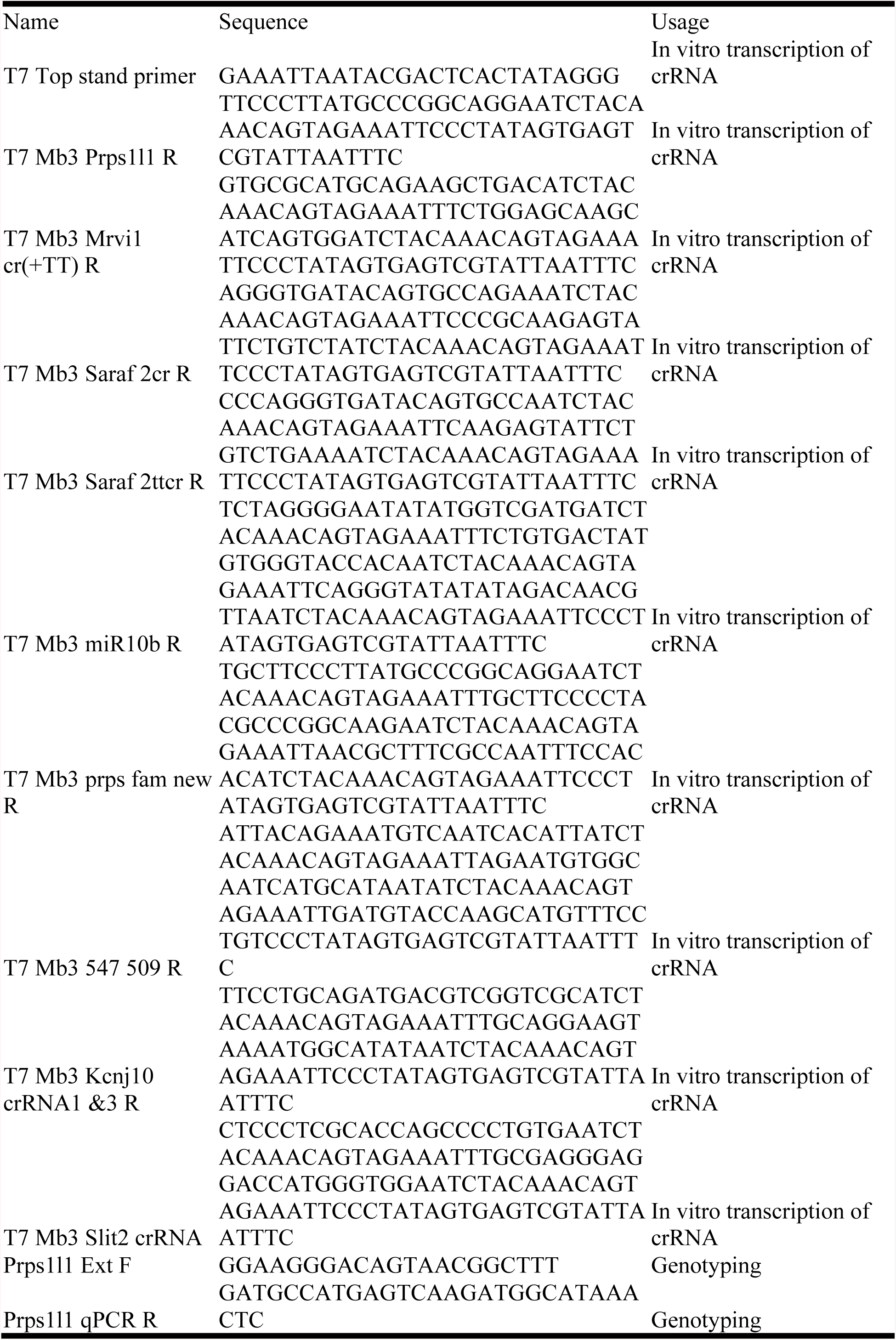
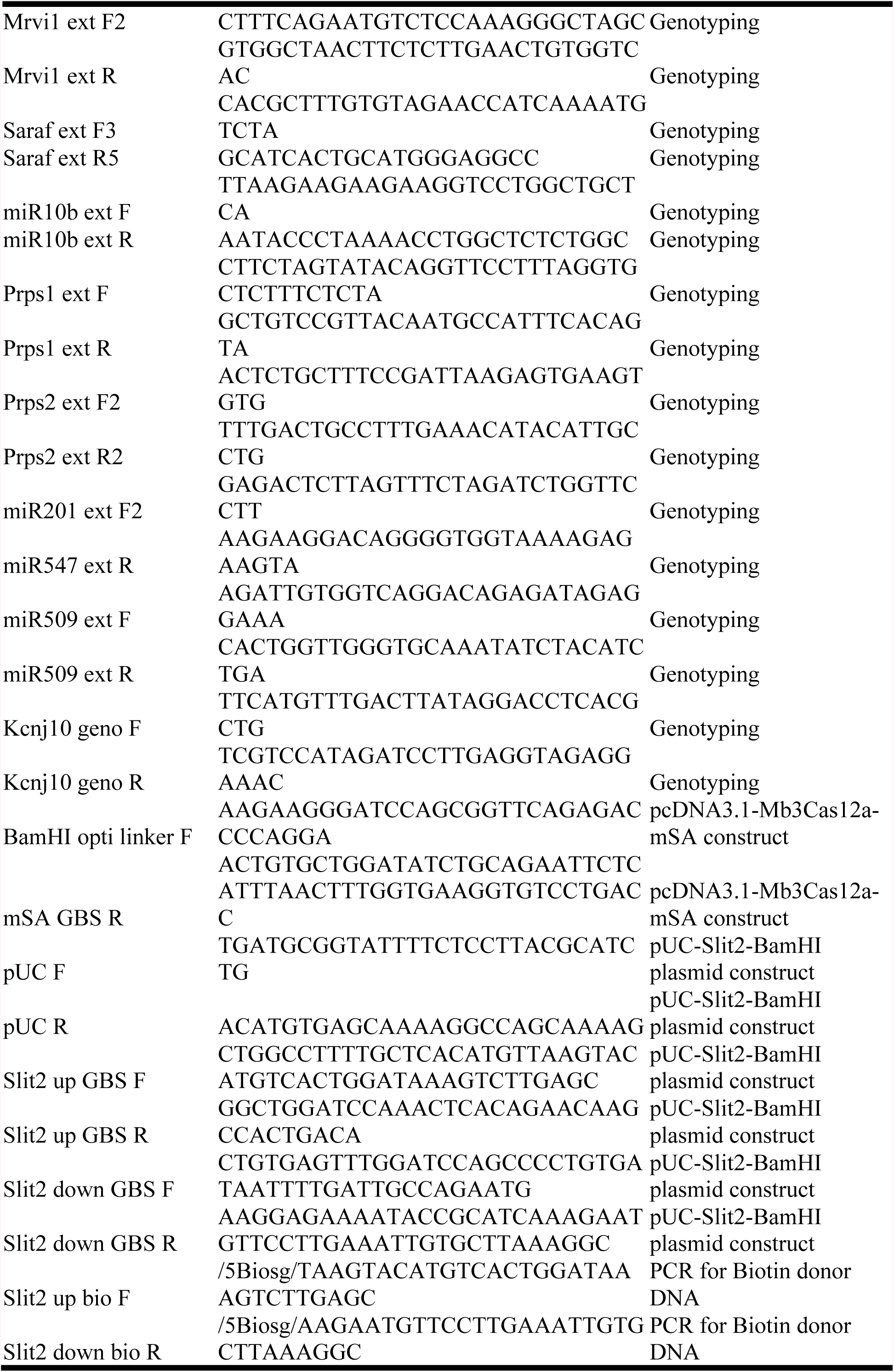
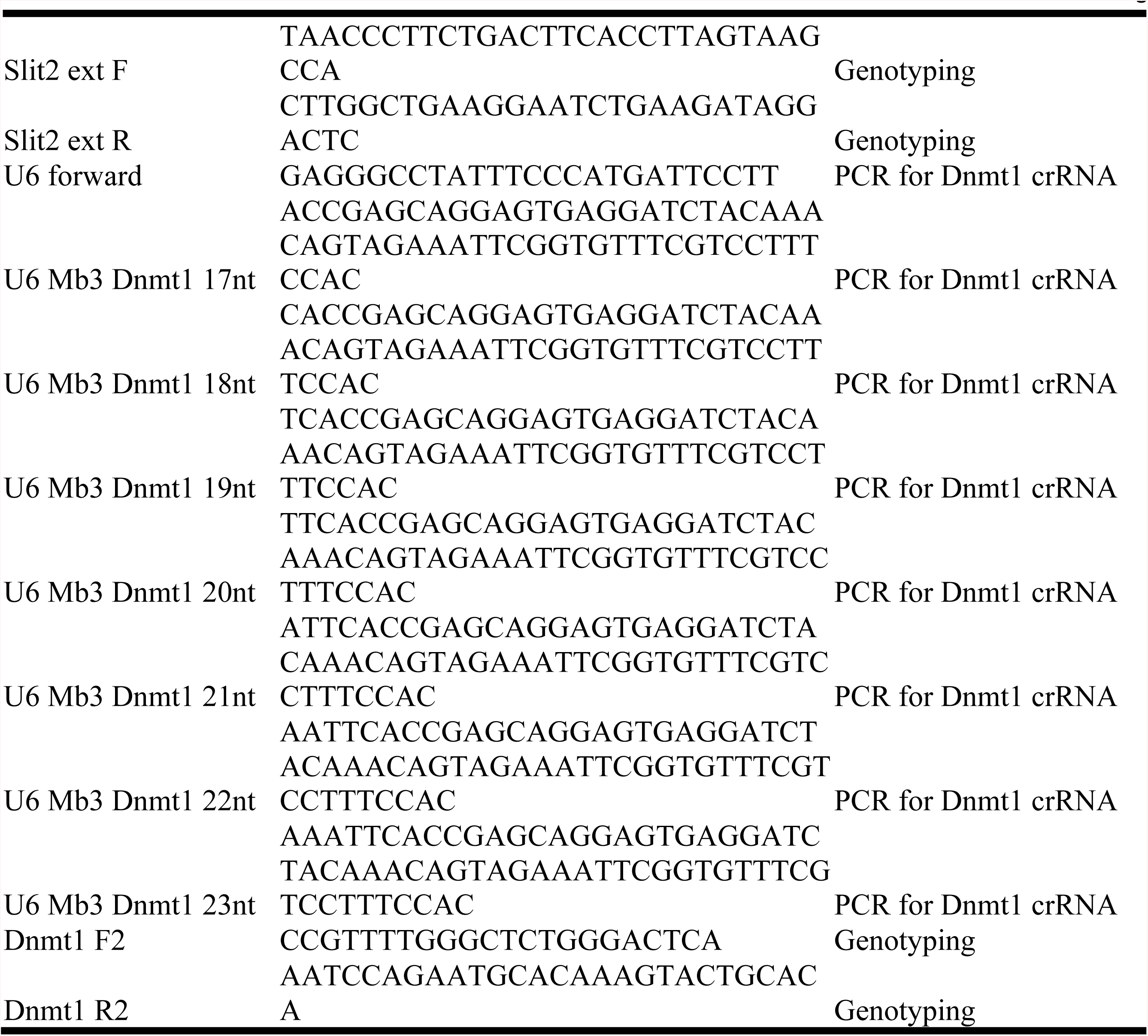
Sequences of primers used in this study.

